# Discoloration of the Cephalothorax in *Pandalus borealis* and its Correlation with Internal Cadmium Concentration

**DOI:** 10.1101/2025.02.02.636125

**Authors:** Nwanneka Joseph, Brandon Reilly, Alex Wight, Xin Yu Huang

## Abstract

This study investigates the correlation between discoloration of the cephalothorax in *Pandalus borealis* (*P. borealis*) shrimp and cadmium concentration. The data, obtained through Atomic Absorption Spectroscopy (AAS), was converted into weight-based cadmium concentrations and analyzed using both parametric and non-parametric statistical tests. Despite assumptions of normal distribution not being met, a two-sample Student’s t-test and Mann-Whitney test were performed to examine the differences in cadmium concentrations between pink and discolored shrimp. Results indicated no statistically significant differences between the two groups. Experimental analysis revealed considerable overlap in cadmium concentration ranges for both groups, with high standard deviations. Factors such as the period of exposure to cadmium and potential influences from other contaminants or microorganisms were also considered. Previous studies have linked discoloration in shrimp to heavy metals and infections, but this study found no correlation between cadmium concentration and discoloration in *P. borealis*. The conclusion suggests that cadmium may not be a sole factor in shrimp discoloration, and other variables should be explored. Recommendations for future research include examining other trace metals, microbial infections, shrimp age, and extent of exposure to provide a comprehensive understanding of cephalothorax discoloration in shrimp.

## I. INTRODUCTION

Wild species of *Pandalus borealis* are found in the icy Northern Atlantic and Arctic oceans where they live 20 to 1330 meters in the ocean (FAO, 2105). The growth phases are slow and they have been known to live for more than 8 years. Instances of discoloration of the cephalothorax have been known to vary seasonally. It is thought that changing feeding patterns and spawning cycles contribute to the discoloration (CAPP, 2015). It is also thought that discoloration of the heads of *P. borealis* indicate heavy metal contamination.

Heavy metals are an intrinsic part of the marine environment. Cadmium being a heavy metal is thought to be generally present in small amounts in the marine environment.

Anthropogenic factors also contribute heavily to cadmium concentration in the environment. Industrial activities have raised concerns and have been linked to contributing immensely to heavy metal contamination. Biota inhabiting sites of contamination are exposed to pollutants. Adverse health effects may result from the ingestion of contaminated foods (Ani, Uhuo, & Nzenwa, 2011). Humans may become contaminated by eating polluted *P. borealis* as well as any other affected aquatic organisms. Aderinola *et al*., (2009) opines that “the capacity of some aquatic organisms to take up heavy metals is up to 10 times the concentration present in the water.” This makes humans especially susceptible in the food chain, especially when such contaminated foods are improperly handled (Ani, Uhuo, & Nzenwa, 2011)

### 1.1. Purpose

Heavy metals are used in many industrial products. Often, they leech into inland water estuaries or directly into the oceans where they can be deposited in sedimentary basins and bioaccumulate in living organisms. (Fleisher *et al*., 1974). A percentage of the deposited metals are released back into open waters over time. Cadmium in particular, is known to be a common environmental pollutant. This project analyzes the bioaccumulation of cadmium in the cephalothorax of *P. borealis* using atomic absorption spectrophotometry. This determined if there was a correlation between discoloration of the carapace and by extension the cephalothorax, and cadmium deposits.

### 1.2. Background

The presence of cadmium in the Canadian marine environment has been attributed to natural and human activities. Being a ubiquitous heavy metal, the natural sources of cadmium to the environment are through diverse methods. Environment Canada (2015) states, “weathering and erosion of cadmium-bearing rocks represent perhaps the most important source.

Approximately 1963 tons of refined cadmium are produced, 23 tons imported, and 1580 tons exported each year in Canada (1992 estimates)”. Other ways that cadmium leaches into the environment are solid waste disposal, and sewage sludge application. Health Canada estimates that 8% of cadmium introduced into the Canadian environment are released on waters. Their reports states that “The most recent estimates identified indicate that base metal smelting and refining operations account for 82% (130 tons) of the total releases to air and water.”

Some studies suggest there is a link between high levels of cadmium present in shrimp and a discoloration of the carapace. Sogegianto, *et al*., (1999) states, “The gills were blackened in shrimps exposed to 2000 and 4000g Cd.L-1. The blackening intensity was higher in shrimp exposed to 4000g Cd.L-1 than those exposed to 2000 g Cd.L-1.” Sogegianto *et al*., (1999) then proceed to state that “the cadmium concentration increased in different tissues (mainly gills, epipodites and hepatopancrease). According to Kargin *et. al*., (2001), “it has been observed that in a significant number of pink shrimps exposed to approximately 760g/ liter (as CdCl2) for an additional nine days in flowing seawater, an unusual blackening of gills occurred.”

The health effects of cadmium were first studied in 1919. Subsequently various health defects became associated with exposure to cadmium including bone disease with fractures, the *itai-itai* disease, and a form of cadmium induced renal osteomalacia. It is important to note the sources of these exposures are mainly through contaminated food and water, and to a lesser extent poor sanitation (Ani, Uhuo, & Nzenwa, 2011). By the 1970s international warnings of health risks from cadmium pollution were issued. Nordberg (2009) states, “The World Health Organization in its International Program on Chemical Safety, WHO/IPCS (1992) identified renal dysfunction as a critical effect. In the 1990’s and 2000 several epidemiological studies have reported adverse health effects, sometimes at low environmental exposures to cadmium in population groups in Japan, China, Europe and USA (reviewed in other contributions to the present volume).”

Various methods of analysis can be used to determine the content of heavy metals in organic samples. Atomic Absorption Spectrometry was originally used as an analytic technique in 1919 by German professors. It is one of the oldest and most reliable forms of quantitative determination of chemical elements. This method uses absorption of light of intrinsic wavelengths by atoms. All atoms are categorized into those having low energies (ground state) and high energies (excited state). Atoms at the ground state absorb energy and move to the excited state. The energy is measured in eV or electron volts and corresponds to the energy of light at a particular wavelength. The atomic density of an atom determines the rate at which it absorbs light. This absorbance is measured for samples containing different concentrations of the element to be tested and plotted on a calibration curve.

### 1.3. Scope

The aim of the project is to determine if there is a link between cadmium concentration in *P. borealis* and the blackening of the cephalothorax. Heavy concentration of cadmium in shrimp has been a proven health risk for decades now and more recent studies have shown that there is a strong correlation of cadmium levels in water and blackening and necrosis of the cephalothorax contained in the carapace. The project specifically makes use of normal pink *P. borealis*, and blackened carapace of *P. borealis* to test for cadmium levels in the samples. The data collected is then passed through a statistical test to determine if there is any statistical correlation.

Shrimp samples were between 4 and 6 cm in length; from the front of the head to the curve of the tail. They were categorically divided into either pink or black based on the corresponding head color. Once categorized, the shrimp heads are removed from the body and added to a composite sample containing a total of 10 shrimp heads of the same category. 62 composite samples were used; 31 from each category. These samples are digested via the wet ash method using aqua regia (3:1 mixture of hydrochloric acid and nitric acid). Following digestion, the samples are filtered and ran through the atomic absorption spectrometer to determine the concentration of cadmium. The data collected is used in a statistical test to determine if there is a statistically significant relationship between the blackening of gills and internal concentration of cadmium.

## II. LITERATURE REVIEW

### 2.1 Cadmium in the Environment

Cadmium is an element which is found naturally in the environment. It is found in many rocks and minerals at the earth crust at relatively low concentrations. There are particular minerals which are known to hold more cadmium. Marine phosphorites for example have been shown to contain 25ppm of cadmium (Fleischer *et al*., 1974). Areas with higher amounts of organic matter and oceanic sediments also have been shown to have relatively high amounts of cadmium. Fresh water usually contains around 1 microgram/ liter, which is higher than the sea averages of about 0.15 micrograms a liter (Fleischer *et al*., 1974).

Although cadmium is shown to be natural in the environment, there are many sources of contamination caused by man. Sewage has been shown to contain large amounts of cadmium. Leaching from landfills has been associated with cadmium contamination (Fleischer *et al*., 1974), (Tyler et al 1989). There also industrial concerns where some factories and mining operations; mostly zinc mining, have been connected to higher cadmium concentrations in the surrounding environment. (Engel and Fowler, 1979). Cadmium is much more easily mobilized in acidic conditions (Tyler *et al*., 1989). Therefore, even pollutions that doesn’t contain cadmium can still increase the cadmium concentrations in the soil and water simply by increasing the acidity, causing leaching to occur from the stones and minerals in the environment (Tyler *et al*., 1989). This means that acid rain is a large concern for cadmium increases to happen in nature. Having this metal in the water and soil is a concern from an environmental perspective and it poses a health issue. Accumulation of cadmium can occur in plants and animals that reside in areas with contaminated soil and water. (Satarug *et al*., 2009).

### 2.2 Health Concerns of Cadmium

Cadmium can be toxic when inhaled or orally ingested, as such this research is focused on the food safety aspects of cadmium and particularly on oral exposures. Ingestions of food contaminants is a source of diseases (Ani, Uhuo, & Nzenwa, 2011), and cadmium has long been known to be toxic, and has been shown to have both chronic and acute symptoms. Acute responses are severe nausea, salivation, vomiting diarrhea, abdominal pains and myalgia (Arena 1963). This can develop into more serious symptoms such as respiratory problems, liver and kidney failure (Flick *et al*., 1971). Death can occur from ingestion of cadmium when consumed at high doses.

Chronic exposure to contaminants (Ani, Uhuo, & Nzenwa, 2011) has been linked to many health issues. It has been shown to build up in the kidney which can have health effects such as kidney diseases including; proteinuria, glomerular filtration problems, and increases the chance of kidney stone formation (ATSDR *et al*.,1997). Reference doses for cadmium were determined from long span studies for both water intake and food consumption. For drinking water, the reference dose is .001 mg/kg/day, and dietary dose is 0.001 mg/kg/day (ATSDR *et al*.,1997).

The best-known example of cadmium toxicity to humans occurred in the 1800s when Japanese people living near a river were all exposed to high levels of cadmium due to a marine nearby (Flick *et al*., 1969). This caused the people to develop a severe disease which was called *Itai-Itai* disease (Edwards and Prozialeck, 2009). This disease was characterized by the softening of bones to the point they would break easily (Flick *et al*., 1969). These symptoms were as a result of several factors.

Cadmium effects the metabolism of vitamin D in the kidney, impairs intake in the gut, and deregulation of collagen metabolism (Nordberg et al., 2007). These factors all affect bone strength and connective tissue strength; which explains the symptoms associated with *Itai-Itai* disease.

Cadmium has also been linked to diabetes. It has been shown that cadmium increases insulin release in the pancreas and can damage the islet of Langerhans cells in the pancreas (Edwards and Prozialeck, 2009).

### 2.3. Shrimp Discoloration

Shrimp are thought to be affected by cadmium concentrations. Soegianto *et al*., *(1998)* performed a study where non-larval shrimp (Penaeus japonicas) were put in controlled environments of different cadmium levels. The cadmium concentrations of the water samples were 0 µg/L, 200 µg/L, 2000 µg/L, and 4000 µg/L. The effects of cadmium on *P. japonicas* and their accumulation location were tested. In lower concentrations the gill color stayed similar to the controls, which was pink. The shrimp exposed to higher concentrations of cadmium (2000µg/L and 4000µg/L) had blackened gills. They attributed this to haemolymphatic vessels being restricted leading to the cells undergoing necrosis.

It was also noted in this study that high levels of cadmium formed in spaces in between the cuticle and epithelial cells, making it prone to bacterial invasion. (Soegianto *et al*., 1998). This suggests a correlation between cadmium concentration in the environment and the blackening of the carapace. The results shown in Figure 2 exposed the locations where higher concentration of cadmium occurred.

**Figure 1:**
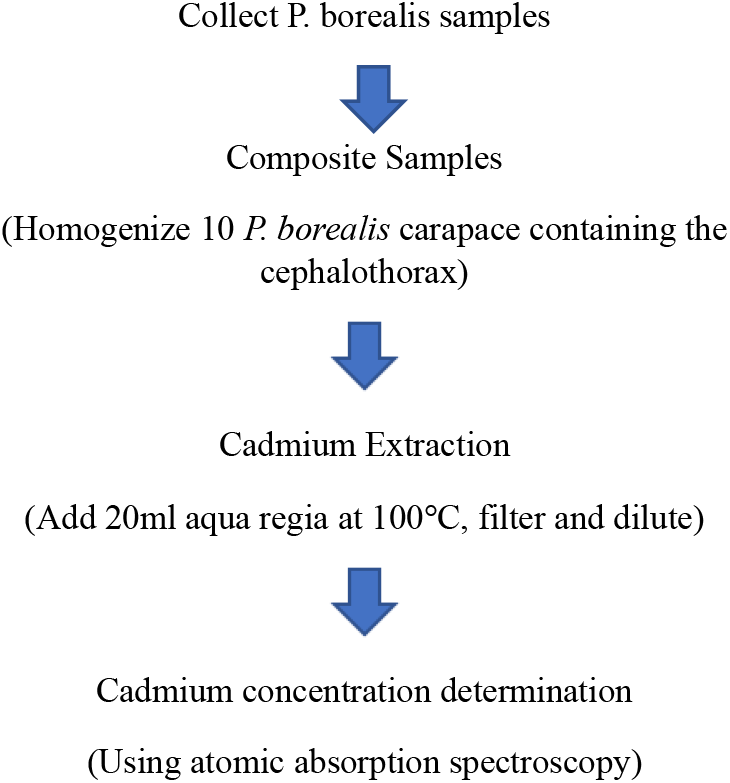
Experimental Procedure

**Figure 2:**
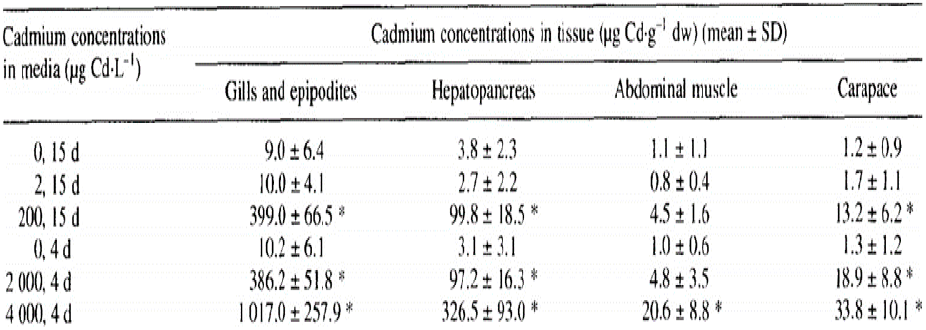
The cadmium concentrations in different organs/areas of the shrimp. Areas tested was the Gills and Epipodites, Hepatopancreas, Abdominal Muscle and Carapace. Image was taken from Soegianto et al., 1998.

Figure 3 shows a clear accumulation of cadmium in the gills, epipodites and hepatopancreas. Both these organs are found in the head area of the shrimp and for simplicity you could say that cadmium accumulates in the head area of the shrimp minus the carapace.

**Figure 3:**
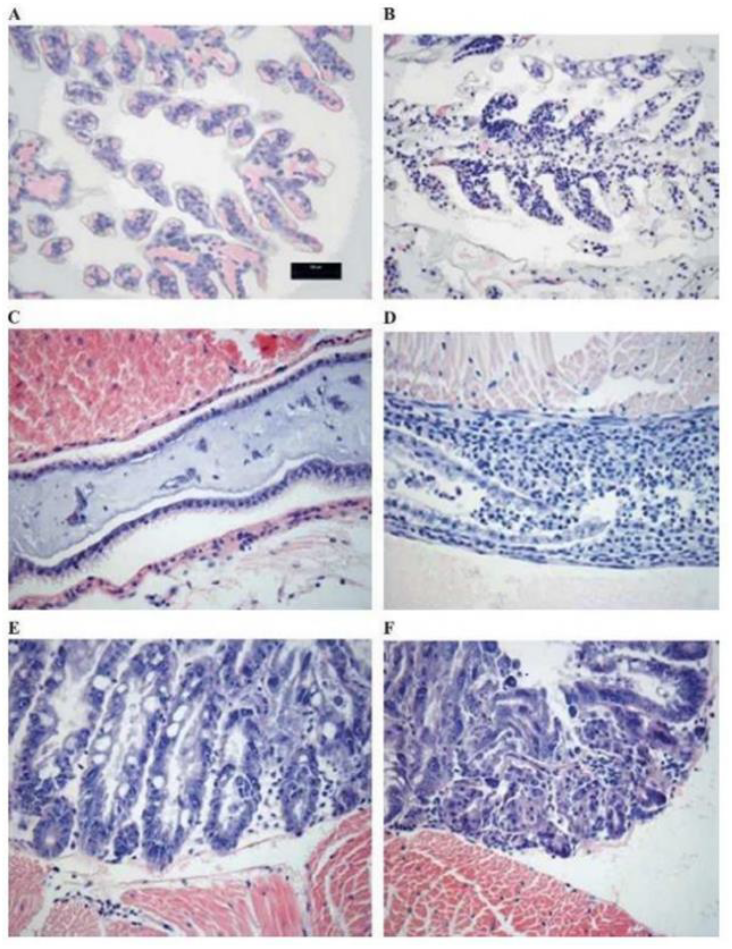
Histological slides from several different organs. A) The gills of a control sample. B) A cadmium exposed sample in which a fair amount of necrosis and sloughing has occurred. C) Picture of the midgut control. D) midgut exposed to cadmium. There was heightened inflammation, liminal debris and ulceration on the cadmium exposed sample. E) Control hepatopancreas. F) Hepatopancreas which has necrosis and fragmentation of tubules. Taken from Keating et al., 2007.

**Figure 4:**
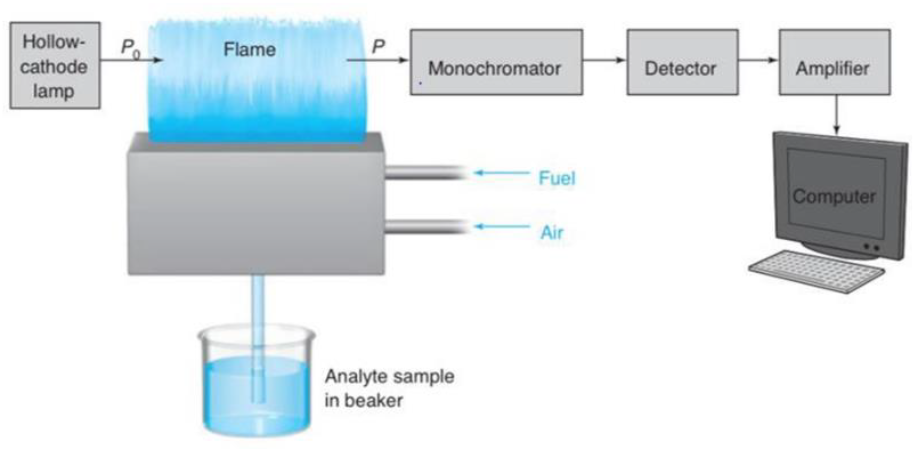
A diagram of the atomic absorption spectrophotometer. Taken from (Harris, D. 2010).

Another study looking at similar experiment was Keating et al., 2007. They used the same principal idea exhibited by Soegianto et al. (1998). Several Litopenaeus vannamei were grown in different concentration of cadmium. Histological studies were carried out comparing the control sample gills (image A) to a shrimp exposed to 10 ppm of cadmium for 28 hours. Necrosis of the gills was observed on the non-control samples. Sloughing also occurred in addition to the discoloration of the gills in this samples (Keating et al., 2007).

The midgut was also examined, as it was considered to be a sensitive organ to stress. The results showed severe (see Image D) (Keating *et al*., 2007). There was damage to the hepatopancreas, which the researchers claimed shows a fair amount of necrosis and sloughing.

Cadmium concentrations in both the studies by Keating *et al*., 2007 and Soegianto *et al*., 1999, have associated environmental cadmium concentration with necrosis and lesions which caused the blackening of the cephalothorax. Both studies used different species of shrimp when performing the experiment and had similar results. Although an experiment has not been conducted on *P. borealis*, it is likely to have similar results.

### 2.4. Other Reasons for Discolorations

There have been additional speculations as to why the cephalothorax of shrimp may become discolored. Trace metals have been associated with the discoloration. Some studies have shown to that copper mat bee a culprit in blackening around the gill region (Lodhi *et al*., 2006). Some authors believe that zooplankton contained in the gut are responsible for the discoloration (Canadian association of Prawn Producers, 2015). This in turn may alter the color of the cephalothorax of the shrimp after harvesting. Females will hold eggs near the gill area which can present as black/green color around the head region (Canadian Association of Prawn Producers, 2015). Seasonal patterns may also affect the color of shrimp, which suggests feeding patterns or spawning cycles could also have an effect on the head color (Canadian association of Prawn Producers, 2015). There may be several reasons the head area of the shrimp may be discolored besides cadmium; however, cadmium has been studied to effect darkening in the head.

### 2.5. Atomic Absorption Spectrophotometry

Atomic Absorption Spectrophotometry (AAS) is a technique used to determine mineral content in a sample. It is often used for heavy metal determination in foods, and common practices have been well established. Some sample preparation is required to get a proper reading on an AAS machine. This involved the removal of organic matter on the sample. There are two ways to accomplish this. One method uses a muffle furnace, which raises the temperature 400-500 degrees Celsius for 24 hours. This ensures that all organic tissues are burned off from the sample.

Metals in the sample are unaffected by this process. The second method uses boiling acid to digest all the organic tissues, breaking down the organic matter and releasing it as a gas. Several acids have been used to do this; of which aqua regia is an example (Tokalioglu *et al*., 2000). Aqua regia is a mix of hydrochloric acid and Nitric Acid. When heated it has been shown to remove organic material in an effective manner which allows analysis of atomic absorption spectrophotometry (Tokalioglu *et al*., 2000).

Atomic absorption spectrophotometry requires a light source which emits the line spectrum of the element being measured, a device for vaporizing the sample, a monochromator, photoelectric detector and a measuring equipment (Royal Society of Chemistry, 2014). Therefore, if the light source can emit a particular wavelength which is absorbed only by the metal being measured it can help determine the amount of that metal in the sample by reading the amount of light of that wavelength passing through the sample (Royal Society of Chemistry 2014). The light source used is a cathode lamp which is made up of the same metal as the sample being measured, the element in the lamp is then excited to produce the right wavelength of light which would be absorbed by the vaporized sample (Royal Society of Chemistry 2014). To vaporize the sample, the sample is heated by putting it through a flame produced by acetylene and oxygen.

This allows the sample to be vaporized and the light source to pass through it for measurement. When the wavelength from the light source is passed through the sample, a percentage of the wavelength is absorbed by the sample, which is a direct relation to the amount of the metal being measured in the sample. The light that passes through can therefore be put through a monochromator for the specific wavelength and measured (Royal Society of Chemistry 2014). The amount of light which passes through can then be compared to a calibration curve of known samples to determine the concentration of the metal being measured (Royal Society of Chemistry 2014).

## III. EXPERIMENT AND RESULT Collect P. borealis samples

### 3.1. Sample Preparation

Shrimp were selected based on size; between 4 and 6 cm from the curve of the tail to the front of the head. Selected shrimp had their heads removed from the body. All extremities including the outer shell were removed from the head.

The composite samples were grinded as shown in Figure 5 using a ceramic mortar and pestle, weighed, and transferred into separate 250 ml volumetric flasks. 25 ml of aqua regia was added to each flask after which they were placed on a heating plate and left to digest for one hour. The sample weights are summarized in the Section 4.0.

**Figure 5:**
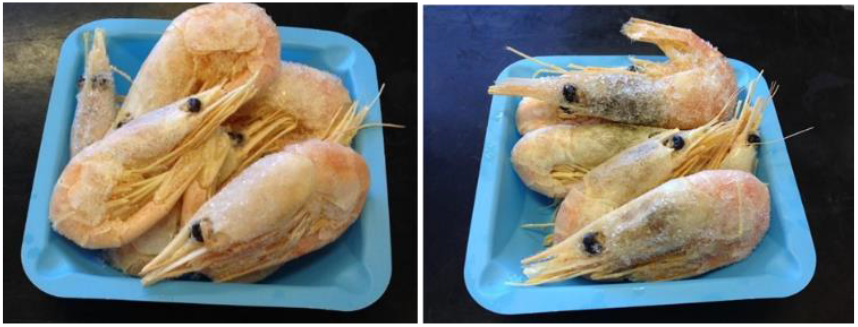
Composite sample of pink headed shrimp (left), and black headed samples (right).

**Figure 6:**
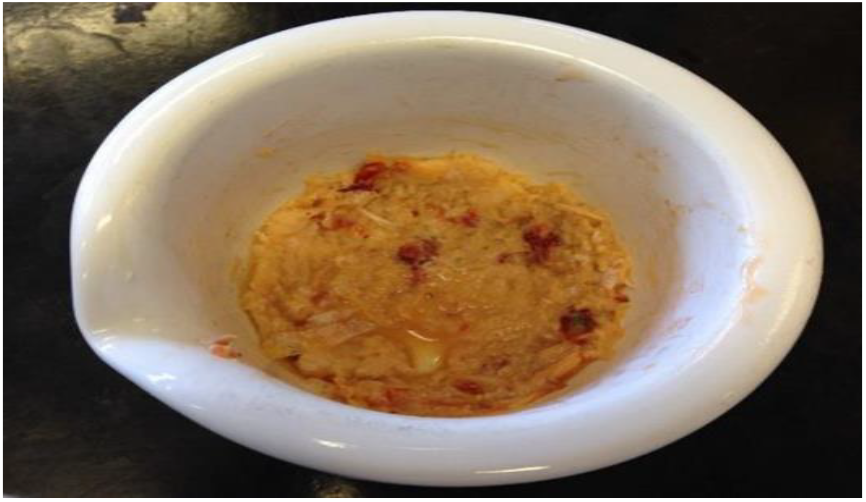
Illustration of a grinded composite sample.

Upon completion of the digestion process, samples were removed from the heating plate and filtered into separate 50ml volumetric flasks. The flasks were then filled to the graduation mark with de-ionized water.

### 3.2 Laboratory Analysis

Instrumental analysis was used to obtain the quantitative data. The particular piece of equipment utilized was the Flame Atomic Absorption Spectrometer (AAS). A set of 5 cadmium standards (0.5, 1, 2, 4, and 8 ppm) with a blank were analyzed prior to sample analysis with an overall R2 of greater than 0.98. Sample data analysis of the obtained values were recorded, and entered in Minitab 17 for statistical analysis. (Refer to Section 4.0).

Atomic Absorption analysis was completed using SHIMADZU AA-6650, accompanied by WizAArd software. This piece of second purge and extinguish was initiated followed by configuration of optic parameters, line search, and beam balance.

### 3.3 Data Analysis

The output from AAS is attached in Appendix B. The cadmium concentrations (mg/L) obtained from the AAS are converted to milligram of cadmium per kilogram of shrimp using equation: equipment required calibration for the proper concentration reading of cadmium, which was completed using element selection. A calibration curve was also created for the 5 experimental standards and the blank prior to sample analysis. Following the entry of experimental parameters in the WizAArd software, additional physical setup of the AAS was required. A 5

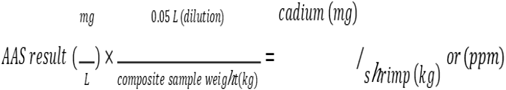

Cadmium concentration in control sample of pink headed shrimp and composite black headed shrimp samples are indicated in the graphical representation below:

### 3.4 Correlation Analysis (Student’s t-Test and Mann-Whitney Test)

To use the student’s t-Test the following assumptions must be met, else the non-parametric Mann-Whitney Test must be used.

#### Mann-Whitney Test

## IV. DISCUSSION

To interpret the collected data, the AAS output were converted into *P. borealis* weight-based cadmium concentrations (mg/kg). The raw data and respective calculations can be found in Section 4.1. The converted data is sorted by date of analysis and are summarized in Table 1-4 of Appendix B. The data is then represented in the Figure 8-9 as bar graphs. In Figure 10, a box plot of the individual concentration of both groups of P. borealis are used to inspect for any outliers. According the Figure 11, no outliers were discovered.

**Table 1:** Summary of the cadmium concentration obtained on January 22, 2015.

| Sample Date | Sample tested (total) | Sample weight (g) | Cadmium Concentration |
| --- | --- | --- | --- |
| 22-Jan-15 | Pink | 4.393 | 6.830 |
| 22-Jan-15 | Pink | 4.334 | 8.844 |
| 22-Jan-15 | Black | 9.7706 | 6.901 |
| 22-Jan-15 | Black | 9.2592 | 6.178 |

**Table 2:** Summary of the cadmium concentration obtained on January 27, 2015.

| Sample Date | Sample tested (total) | Sample weight (g) | Cadmium Concentration |
| --- | --- | --- | --- |
| 27-Jan-15 | Pink | 10.040 | 4.959 |
| 27-Jan-15 | Pink | 10.157 | 5.477 |
| 27-Jan-15 | Pink | 10.282 | 4.650 |
| 27-Jan-15 | Pink | 10.471 | 4.783 |
| 27-Jan-15 | Pink | 10.507 | 2.482 |
| 27-Jan-15 | Pink | 10.260 | 4.036 |
| 27-Jan-15 | Pink | 10.387 | 5.058 |
| 27-Jan-15 | Pink | 10.122 | 4.901 |
| 27-Jan-15 | Pink | 10.343 | 4.203 |
| 27-Jan-15 | Black | 10.114 | 5.580 |
| 27-Jan-15 | Black | 10.772 | 5.355 |
| 27-Jan-15 | Black | 10.772 | 5.387 |
| 27-Jan-15 | Black | 10.084 | 6.036 |
| 27-Jan-15 | Black | 11.218 | 5.675 |
| 27-Jan-15 | Black | 10.388 | 6.442 |
| 27-Jan-15 | Black | 10.353 | 6.352 |
| 27-Jan-15 | Black | 10.126 | 6.567 |
| 27-Jan-15 | Black | 10.994 | 6.548 |

**Table 3:** Summary of the cadmium concentration obtained on February 3, 2015.

| Sample Date | Sample tested (total) | Sample weight (g) | Cadmium Concentration |
| --- | --- | --- | --- |
| 3-Feb-15 | Pink | 10.278 | 5.451 |
| 3-Feb-15 | Pink | 10.134 | 4.254 |
| 3-Feb-15 | Pink | 10.003 | 7.029 |
| 3-Feb-15 | Pink | 10.292 | 5.201 |
| 3-Feb-15 | Pink | 10.043 | 5.448 |
| 3-Feb-15 | Pink | 10.039 | 5.451 |
| 3-Feb-15 | Pink | 10.868 | 4.828 |
| 3-Feb-15 | Pink | 10.369 | 4.394 |
| 3-Feb-15 | Pink | 10.028 | 3.956 |
| 3-Feb-15 | Pink | 10.433 | 5.968 |
| 3-Feb-15 | Black | 10.323 | 3.851 |
| 3-Feb-15 | Black | 10.028 | 2.405 |
| 3-Feb-15 | Black | 10.437 | 4.410 |
| 3-Feb-15 | Black | 10.195 | 4.149 |
| 3-Feb-15 | Black | 10.298 | 4.071 |
| 3-Feb-15 | Black | 10.219 | 4.002 |
| 3-Feb-15 | Black | 10.138 | 3.420 |
| 3-Feb-15 | Black | 10.061 | 4.689 |
| 3-Feb-15 | Black | 10.184 | 5.269 |
| 3-Feb-15 | Black | 10.407 | 4.696 |

**Table 4:** Summary of the cadmium concentration obtained on February 10, 2015.

| Sample Date | Sample tested (total) | Sample weight (g) | Cadmium Concentration |
| --- | --- | --- | --- |
| 10-Feb-15 | Pink | 10.673 | 6.629 |
| 10-Feb-15 | Pink | 10.225 | 7.831 |
| 10-Feb-15 | Pink | 10.089 | 8.565 |
| 10-Feb-15 | Pink | 10.454 | 7.620 |
| 10-Feb-15 | Pink | 10.145 | 7.833 |
| 10-Feb-15 | Pink | 10.104 | 7.952 |
| 10-Feb-15 | Pink | 10.725 | 5.577 |
| 10-Feb-15 | Pink | 10.050 | 6.597 |
| 10-Feb-15 | Pink | 10.027 | 10.895 |
| 10-Feb-15 | Pink | 10.655 | 9.111 |
| 10-Feb-15 | Black | 10.063 | 5.555 |
| 10-Feb-15 | Black | 10.027 | 6.524 |
| 10-Feb-15 | Black | 10.453 | 5.832 |
| 10-Feb-15 | Black | 10.517 | 4.808 |
| 10-Feb-15 | Black | 10.271 | 5.613 |
| 10-Feb-15 | Black | 10.598 | 6.186 |
| 10-Feb-15 | Black | 10.202 | 6.419 |
| 10-Feb-15 | Black | 10.599 | 5.765 |
| 10-Feb-15 | Black | 10.160 | 6.798 |
| 10-Feb-15 | Black | 10.149 | 6.845 |

**Figure 7:**
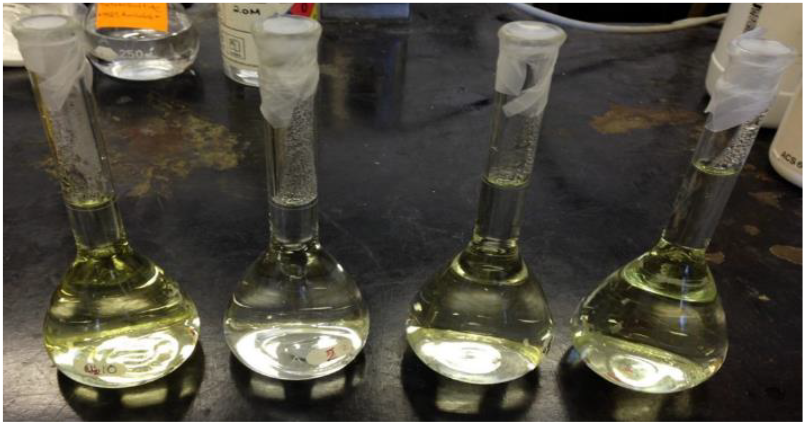
Diluted solution of the composite samples.

**Figure 8:**
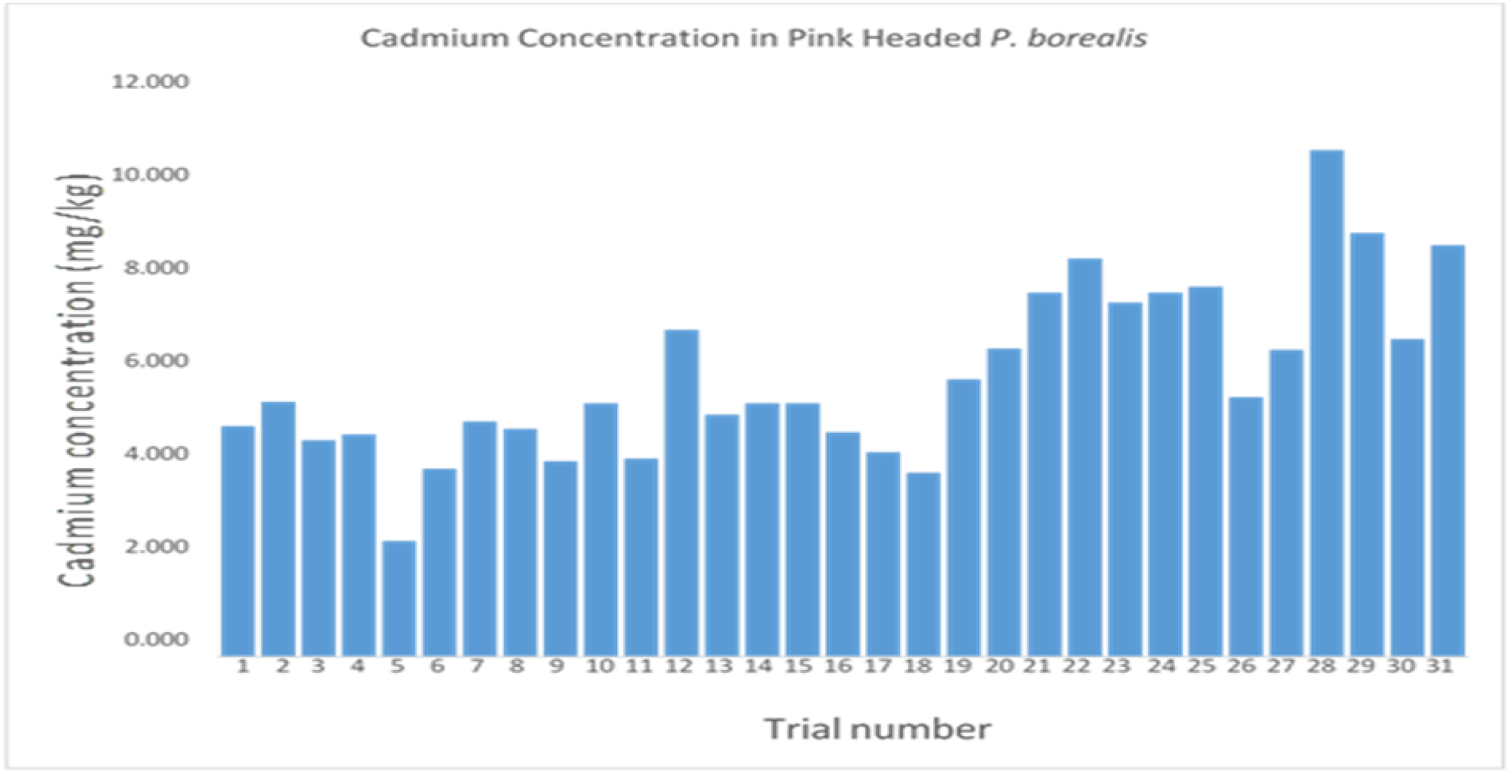
Cadmium concentration measured in the composite sample of pink headed P. borealis

**Figure 9:**
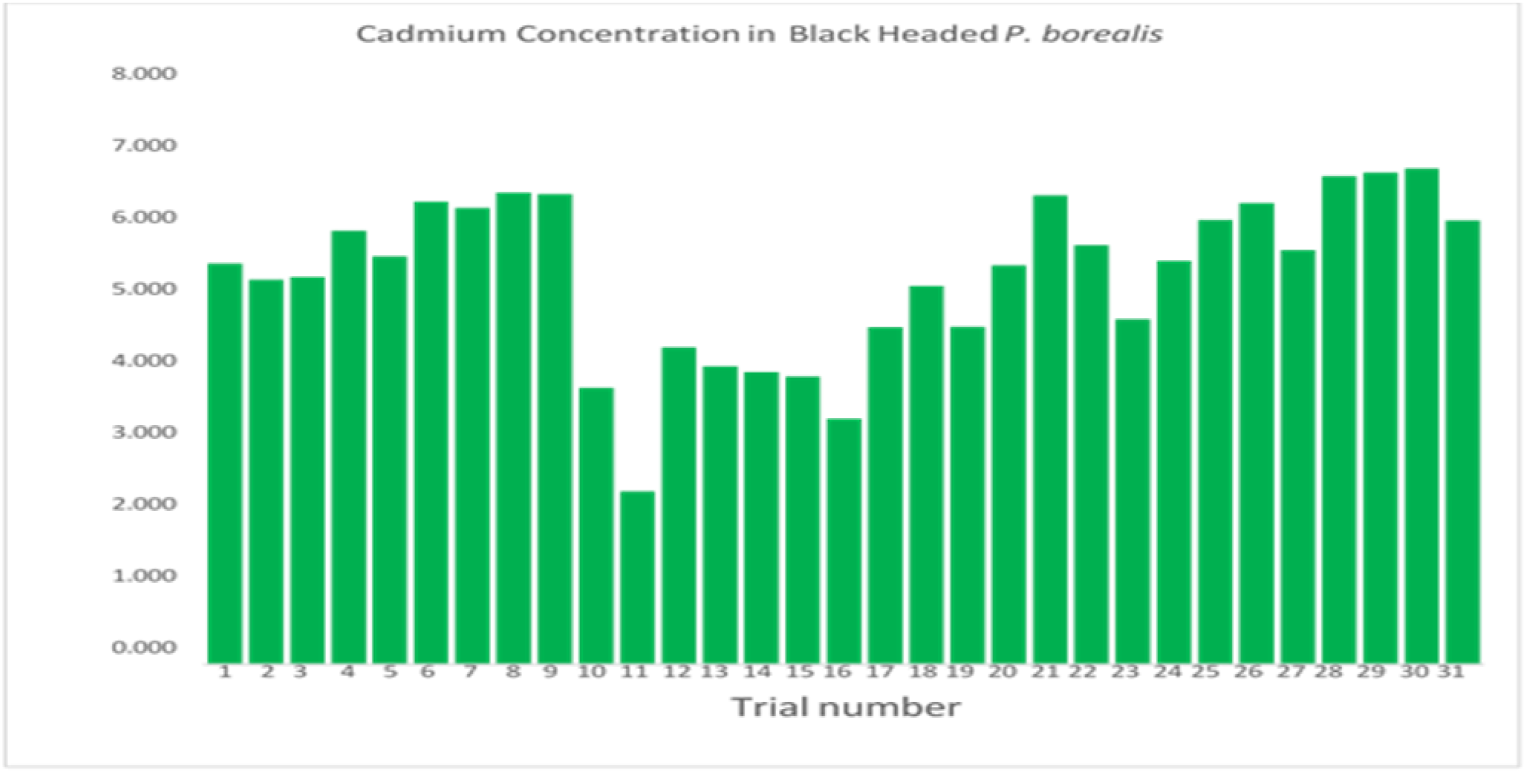
Cadmium concentration measured in the composite sample of black headed P. borealis.

**Figure 10:**
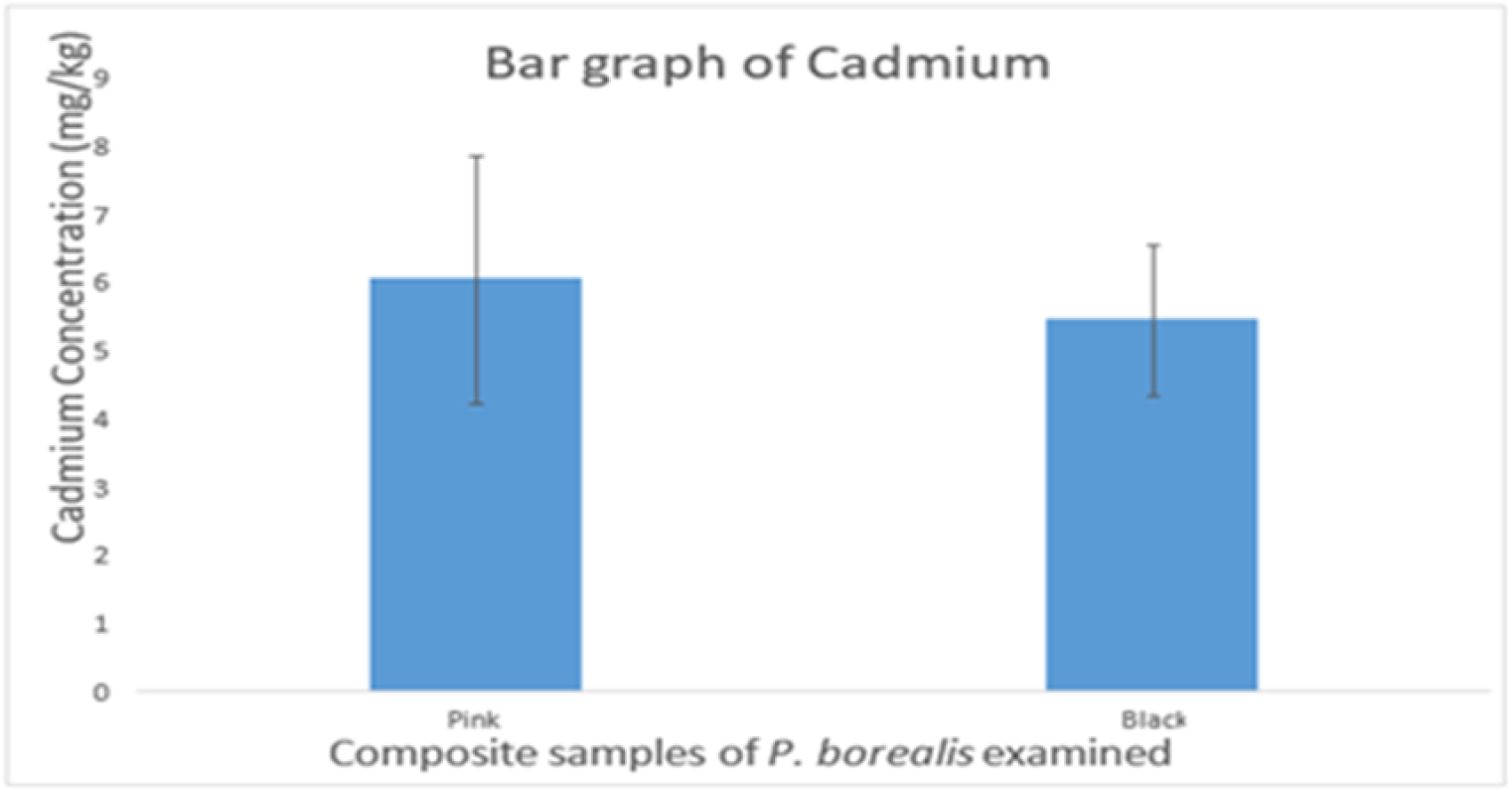
The average cadmium concentration measured in the composite samples of the *P. borealis* examined The error bar represents the standard deviation (Pink = 6.026 ±1.823 mg/kg, Discolored=5.430±1.111 mg kg). The relative standard deviation is (pink - 30 2 5 7%, black - 20 469)

**Figure 11:**
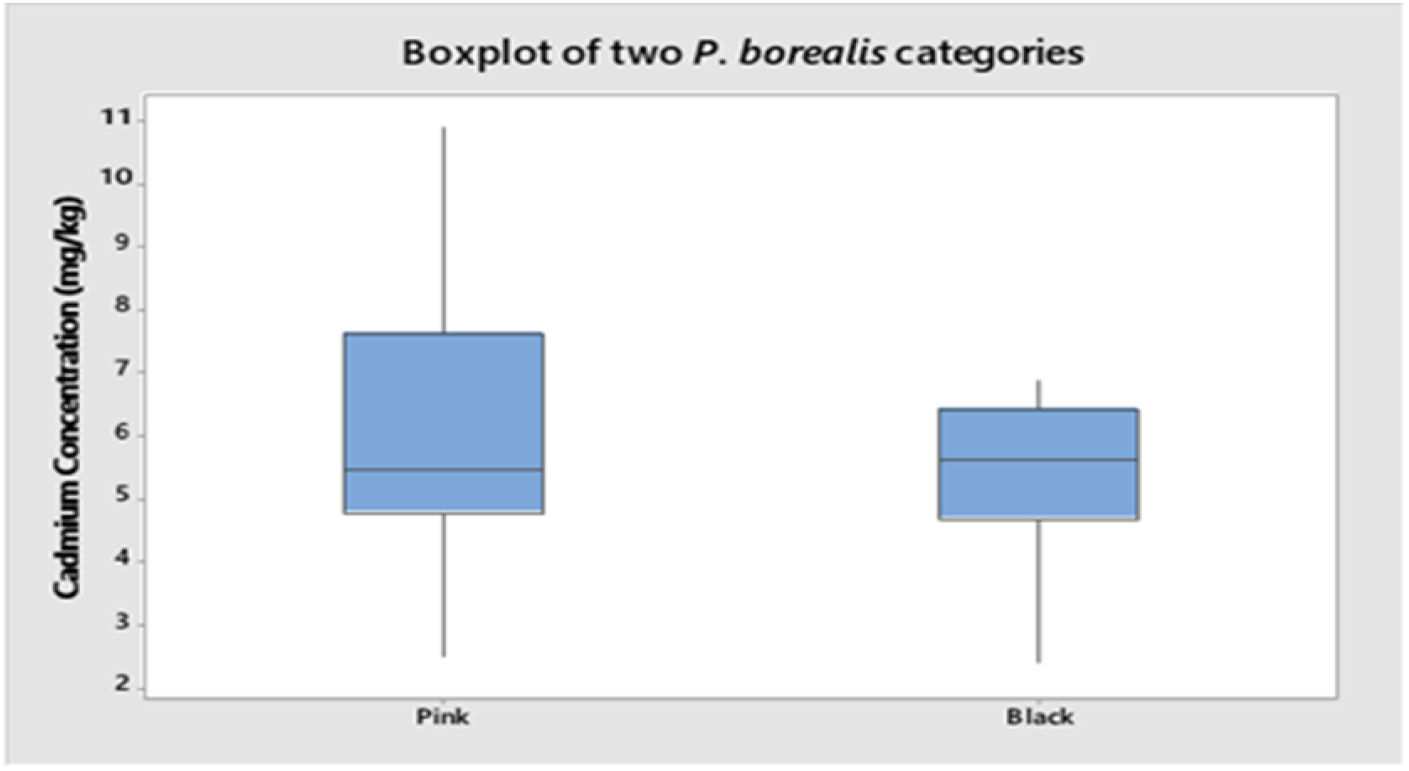
Boxplot is used to visually examine if any outliers exist in the data. No outliers were found.

### 4.1 Correlation Analysis (Student’s t-Test and Mann-Whitney Test)

#### Student’s t-test

To examine the correlation of discoloration of the cephalothorax and cadmium concentration of the two sample, Student’s t-Test was used. The two sample Student’s t-Test is used to determine if there was a significant difference between the mean cadmium concentration in the pink and discolored shrimps. In order to use the t-Test the following assumptions must be met:

1. The cadmium concentration of both of the populations (pink or black) from which the samples were selected is normally distributed or approximately normal.
2. The variances S 1^2^ and S 2^2^ of the two populations are equal.
3. The random samples are selected in an independent manner from the two populations.

The hypothesis for determining if the cadmium concentration of both groups is normally distributed is as follows:

Ho = Normally distributed

Ha = Not normally distributed

α = 0.1

As shown in Figure 12 and Figure 13, concentrations for both groups are not normally distributed, the p-value for pinked headed samples is 0.073, and 0.088 for discolored P. borealis samples. Both p-values are less than 0.10, therefore, the null hypothesis that the sample is normally distributed is rejected, and the non-parametric test (Mann-Whitney Test) should be used. Since sample size is greater than 30, the t-test was conducted and the results are shown in Appendix A.

**Figure 12:**
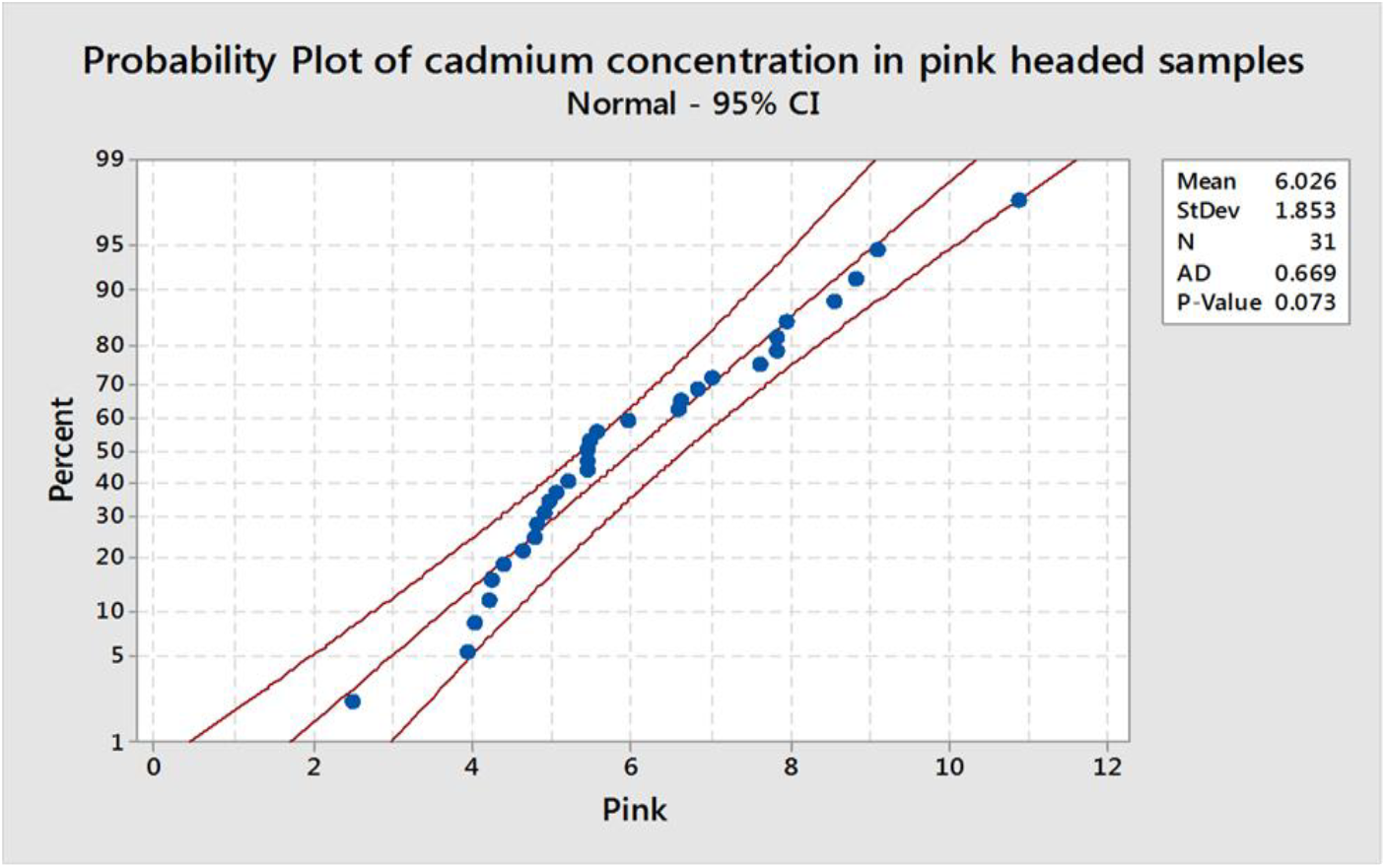
Probability plot of the pink headed composited samples. The p-value is less than 0.1. Therefore, this represents non-normal distribution

**Figure 13:**
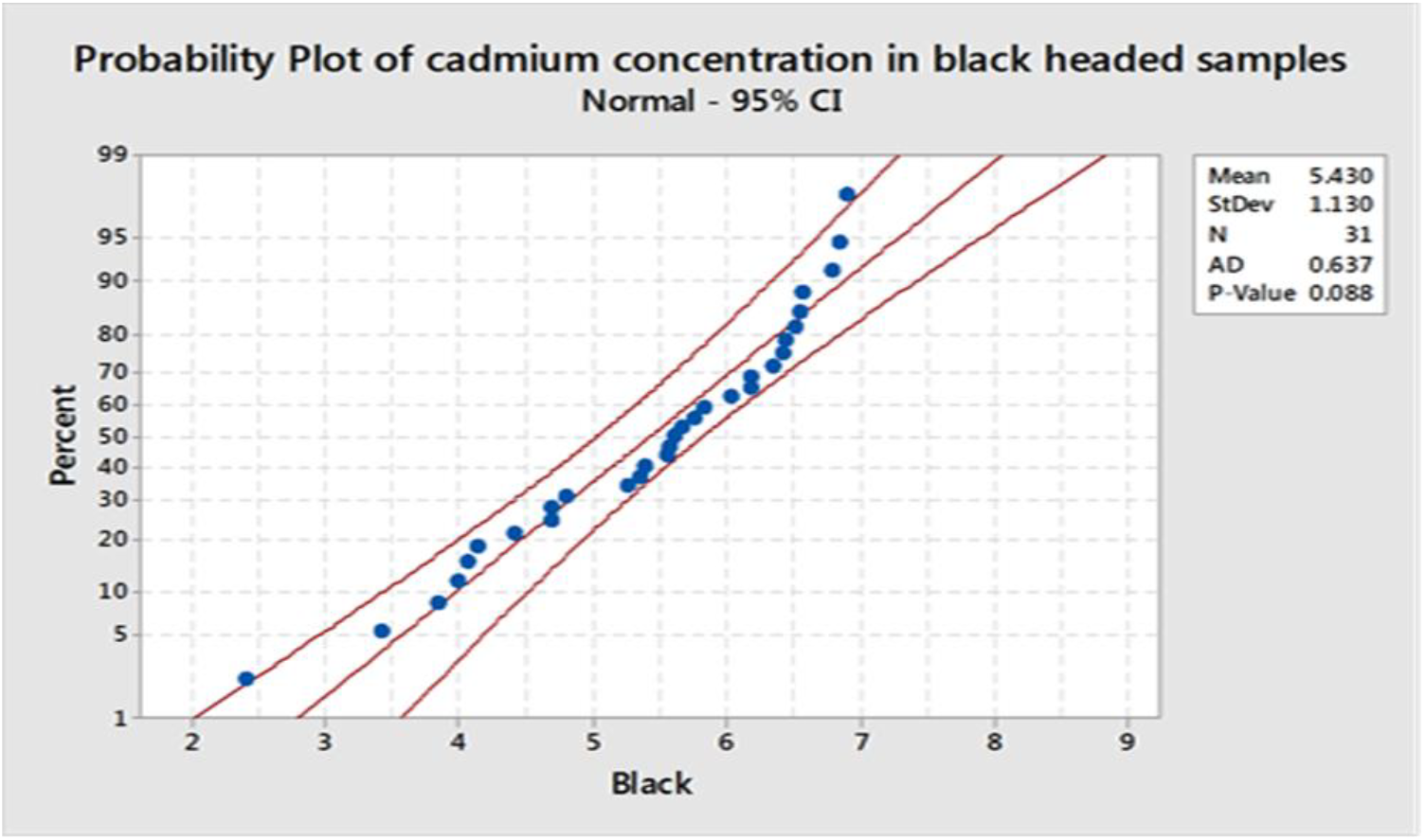
Probability plot of the blacked headed composited samples. The p-value is less than 0.1. Therefore, this represents non-normal distribution

#### Mann-Whitney Test

In order to use the Mann-Whitney to be used the following assumptions must be met:

1. One dependent variable will be measured at the continuous or ordinal level.
2. One independent variable will be measured that consists of two categories, or independent groups.
3. There should be no relationship between the observation in each group of the independent variable or between the groups themselves.
4. Determine if the distribution of concentration for each group of the independent variables have the same or different shape. If the shape is the same, then the Mann-Whitney test can be used to determine if there are equal distributions. If the shape is different, the test is used to determine if the median concentration of both categories is the same.

The data obtained satisfies assumption 1-3, and it does not have the shape distribution (as shown in Figure 8 and 9). Therefore, the Mann-Whitney test is used to determine if median is the same. The hypothesis for determining if the cadmium concentrations of both groups are related using the Mann-Whitney test is as follows:

Ho =, 11pink = 11black (11 is the sample median)

Ha = 11 pink ≠ 11 black

α = 0.05

When the Mann-Whitney test was conducted, the p-value was found to be 0.3983, which is greater than α = 0.05 as shown in Figure 14. Therefore, we failed to reject our null hypothesis and can conclude that there is no significant difference between the cadmium concentration in pink and discolored *P. borealis*.

**Figure 14:**
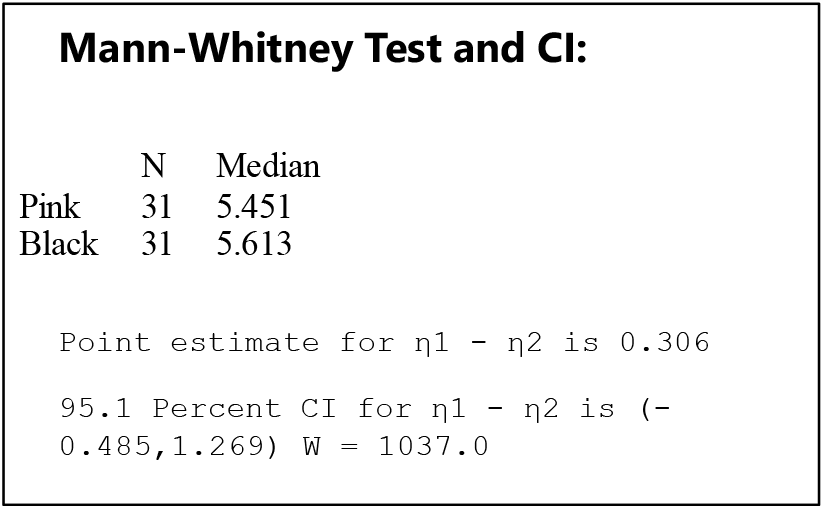
Results obtained from the Mann-Whitney test for two sample mean.

**Figure 15:**
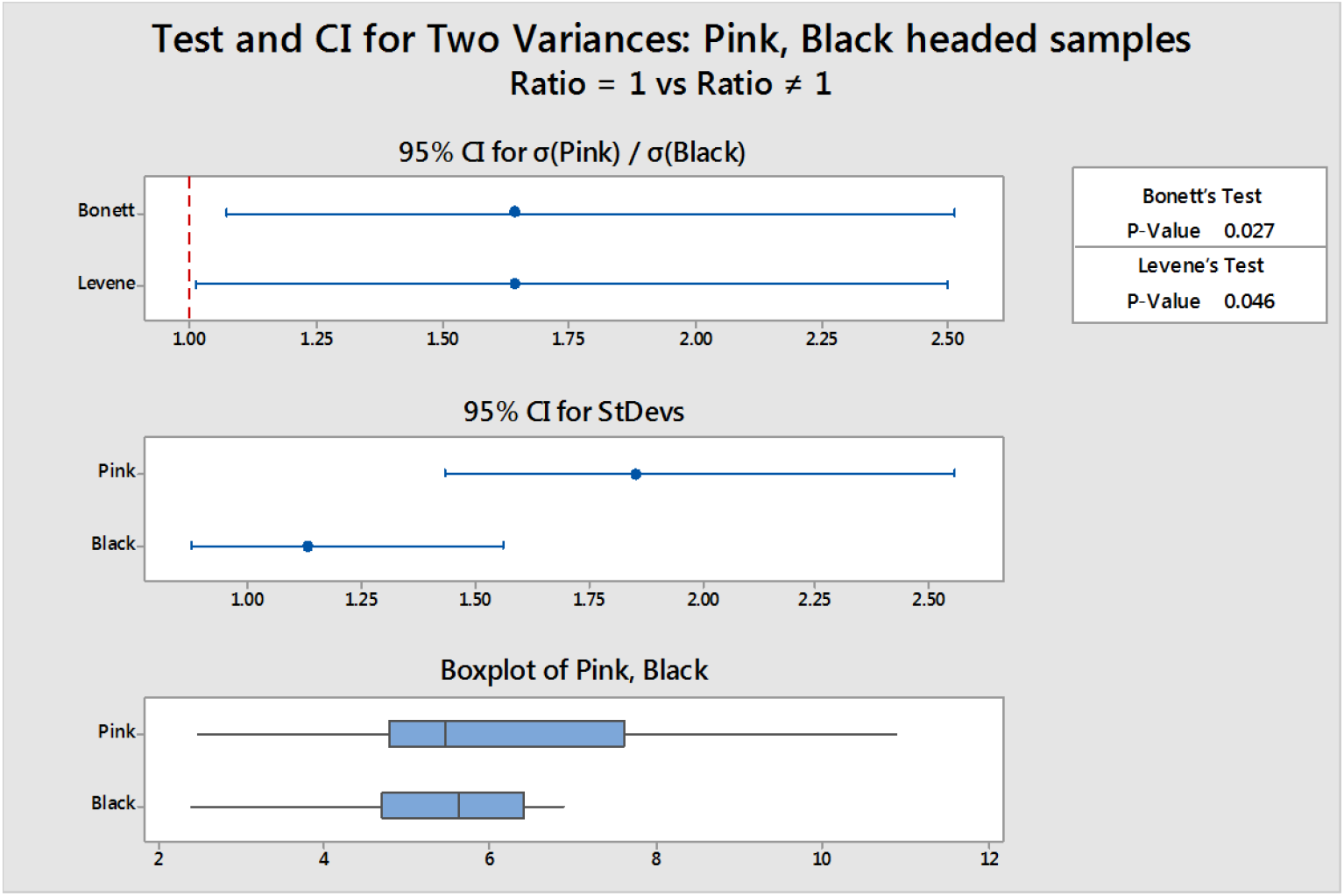
Test for equal variances comparing the ratio of the standard deviations. The p-value in the Leven’s test (0.046) is less than the level of significance (α=0.05), thus the null hypothesis is rejected and the variances are not equal.

**Figure 16:**
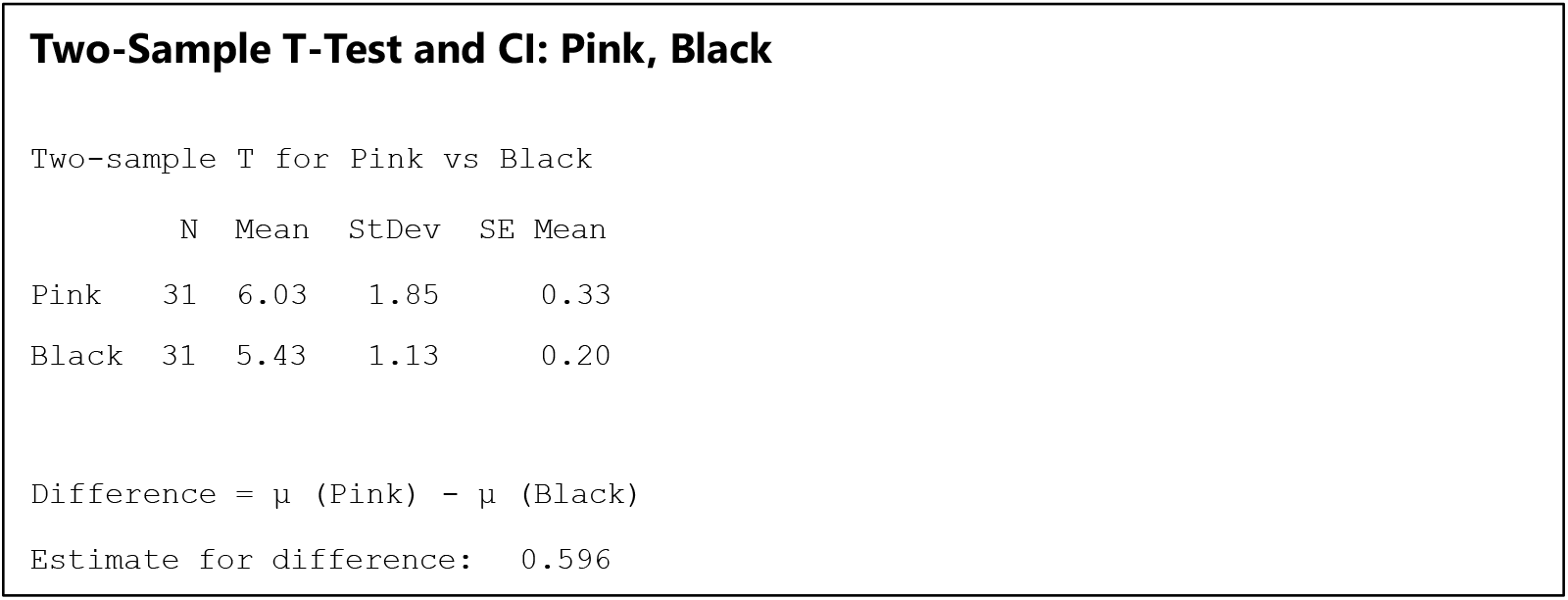
Results obtained from the two-sample t-test test for two samples mean.

### 4.2 Experimental Analysis

As summarized in section 5.1, the results obtained above, there were no statistically significant difference observed. One factor that has affected the results is the large range of concentration obtained for the pink headed P. borealis, as shown in Figure 10. The concentration ranges from 2.484 mg/kg to 10.895 mg/kg for the pink headed *P. borealis*. The range for black headed *P. borealis* ranges from 2.405 to 6.901 mg/kg. The standard deviation is very high for both categories, as the relative standard deviation is 30% for the pink headed samples and 20% for the black headed samples. Thus, there is a considerable amount overlap of the data from both groups. Furthermore, the period of exposure would affect the cadmium concentration, since the longer the shrimp is exposed, the more necrosis can occur. According to Soegianto (1999), exposure at 200 µg/L for 15 days did not display any structural alterations in the gills, while 4-day exposure at 2000 µg/L and 4000 µg/L of cadmium resulted in profound changes.

In previous studies, other factors have been shown to influence the discoloration of the cephalothorax in shrimp such as heavy metals and microorganisms. In the study by Li et al. (2009) on giant freshwater prawn (*Macrobrachium rosenbergii*), exposure to copper has resulted in structural changes in the gills and hepatopancreases. The degree of damage observed in both tissues were related to the elevated copper concentration (Li et al., 2009). According to Soegianto et al. (1999) bacterial infections of the gills contribute to the color and tissue alteration in the shrimp *Penaeus japonicas*. Fungal infection by Fusarium is the cause of black gill disease that is found for *P. Japonicas* (Reza, 2014).

In addition to chemical contaminates or microorganisms, zooplankton may be present in the digestive tract of some shrimps that may cause the head to become brown or black in color (Canadian Association of Prawn Producers, 2015). Also, matured eggs with in the ovary of the shrimp can give green color in the head.

## V. CONCLUSION AND RECOMMENDATIONS

Based on the results from our laboratory and statistical analysis, it is concluded that there is no correlation found between shrimp discoloration and cadmium concentration. Although preceding studies have linked blackening of the gills to a higher cadmium concentration, the data from our study did not support those findings. It is important to note that this experiment focused on cadmium concentrations; there could be many other factors that influence the color of the shrimp cephalothorax.

More expansive studies would need to be conducted to examine other causes for the discoloration of cephalothorax in shrimp. The recommendation is to include a more in-depth analysis of the following:

- Other trace metals present such as copper.
- Impact of microbial infections in the shrimp.
- Age of the shrimp.
- Extent of exposure including the magnitude, duration, and frequency.

## Appendices

## Appendix A Student’s t-Test

## Appendix B

The following pages readings obtained from the atomic absorption spectrometer:

## REFERENCES

[1] Aderinola, O.J, Clarke, E.O, Olarinmoye, O.M, Kusemiju, V and Anatekhai, M.A. (2009). Heavy Metals in Surface Water, Sediments, Fish and Periwinkles of Lagos Lagoon. American-Eurasian J. Agric. & Environ. Sci., 5 (5): 609–617. ISSN 1818-6769 © IDOSI Publications, Lagos.

[2] Ani, O.C., Uhuo, A.C, and Nzenwa, N.J. (2011). The prevalence of Staphylococcus aureus in food samples at selected areas of Aba, Abia State, Nigeria. Continental Journal of Biological Sciences. 4(2): 1–5.

[3] Arena, J. (1963) Poisoning Chemistry-Symptoms-Treatment. Thomas Springfeild 99–127.

[4] Canadian association of prawn producers. (2015). Key facts about cold water shrimp. Cold water shrimp. Retrieved March 18, 2015 from http://www.shrimp-canada.com/coldwater-shrimp/key-facts-about-the-coldwater-shrimp.html.

[5] Edwards, J. and Prozialeck, W. (2009). Cadmium, diabetes and chronic kidney disease. Toxicology and Applied Pharmacology 238: 289–293.

[6] Engel, D. and Fowler B. (1979). Factors Influencing Cadmium Accumulation and Its Toxicity To Marine Organisms. Environmental Health Perspectives. 28:81–88.

[7] Environment Canada. (1994). Health Canada: Cadmium and its compounds. ISBN: 0-662-22046-3. Cat. No.: En 40-215/40E.

[8] Flick, D., Kraybill, H. and Dimitroff, J. (1971). Toxic Effects of Cadmium a Review. Environmental Research 4. 71–85.

[9] Nordberg, G.F., Nogawa, K., Nordberg, M., and Friberg, L. (2007) “Cadmium,” in Chapter 23 in Handbook of the Toxicology of Metals, G. F. Nordberg, B. F. Fowler, M. Nordberg, and L. Friberg, Eds., pp. 445–486, Elsevier, Amsterdam, The Netherlands, 3rd edition.

[10] Harris, D. (2010). Quantitative chemical analysis. New York: W.H. Freeman.

[11] Keating, J., Delaney, M., Meehan-Meola, D., Warren, W., Alcivar, A. and Warren, A. (2007). Histological Findings, Cadmium Bioaccumulation, and Isolation of Expressed Sequence Tags in Cadmium Exposed, Specific Pathogen Free Shrimp, Litopenaeus Vannamei Postlarvae.

[12] Li, N., Zhao, Y., & Yang, J. (2009). Impact of Waterborne Copper on the Structure of Gills and Hepatopancreas and its Impact on the Content of Metallothionein in Juvenile Giant Freshwater Prawn Macrobrachium rosenbergii (Crustacea: Decapoda). Archives Of Environmental Contamination and Toxicology, 56(4), 811–812. doi:10.1007/s00244-009-9306-y.

[13] Lodhi, H., Khan, M., Verma, R. and Sharma, U. (2006). Acute toxicity of copper sulphate to fresh water prawn. Journal of Environmental Biology 27: 585–588.

[14] Moreno, R.P.A., Medesani, D.A. and Rodrıguez, E.M. (2003). Inhibition of molting by cadmium in the crab Chasmagnathus granulate (Decapoda; Brachyura). Aquat.Toxicol., 64: 155–164.

[15] Pierron, F., Baudrimont, M., Boudou, A., and Massabuau, J.C. (May 2007). Effects of salinity and hypoxia on cadmium bioaccumulation in the shrimp Palaemon longirostris. Environ Toxicol Chem. 2007 May; 26(5):1010–7.

[16] Reza, S., Mahdi, P., Hossein, K., Nahid, S., Hamid, B., Saeid, M., & Mohammad, F. (2015). Studying Histopathology of Black Gill Disease in Marine Shrimp of Bandar Abbas Coast. Unique Journal of Pharmaceutical and Biological Sciences, 2(2), 11–15.

[17] Royal Society of Chemistry. (2014). Atomic absorption spectrometry. Retrieved from http://www.kau.edu.sa/Files/130002/Files/6785_AAs.pdf

[18] Salaramoli, J., Salamat, N., Razavilar, V., Najafpour, Sh. and Aliesfahani, T. (2012). A Quantitative Analysis of Lead, Mercury and Cadmium Intake by Three Commercial Aquatics, Hypophthalmichthys molitrix, Onchorhynchus mykiss (Walbaum) and Fenneropenaeus indicus. World Applied Sciences Journal 16 (4): 583–588, 2012 ISSN 1818-4952 © IDOSI Publications.

[19] Shimadzu Corporation. (n.d.). ATOMIC ABSORPTION SPECTROPHOTOMETRY COOKBOOK. Section 2. Retrieved March 19, 2015, from http://blog.utp.edu.co/docenciaedwin/files/2014/03/2Standard-Sample-Preparation-Method.pdf

[20] Tokalioglu, S., Kartal, S., Elci, L. (2000). Determination of heavy metals and their speciation in lake sediments by flame atomic absorption spectrometry after a four-stage sequential extraction procedure. Analytica Chimica ACta 413:22–40

[21] Tyler, G., Balsberg Pahlsson, M., Bengtsson G., Baath, E., Tranvik, L. (1989). Heavy Metal Ecology of Terrestrial Plants, Microorganisms and Invertebrates. Water, Air and Soil Pollutions 47: 189–215.

[22] Wu, J., Chen, H. and Huang, D. (2008). Histopathological and biochemical evidence of hepatopancreatic toxicity caused by cadmium and zinc in the white shrimp, Litopenaeusvannamei. Chemos., 73: 1019–102

[23] Wu, J., Chen, H. and Huang, D. (2008). Histopathological and biochemical evidence of hepatopancreatic toxicity caused by cadmium and zinc in the white shrimp, Litopenaeusvannamei. Chemos., 73: 1019–102.

